# Disentangling the historical routes to community assembly in the global epicentre of biodiversity

**DOI:** 10.1101/2022.05.18.492504

**Authors:** Bouwe Rutger Reijenga, Benjamin G Freeman, David J Murrell, Alex L Pigot

## Abstract

**Aim:** The coexistence and turnover of species along elevation gradients makes tropical mountains hotspots of biodiversity. However, understanding the historical processes through which species arising in geographic isolation (i.e. allopatry) assemble along the same mountain slope (i.e. sympatry) remains a major challenge. Multiple models have been proposed including, (1) the sorting of already elevationally divergent species, (2) the displacement of elevation upon secondary contact, potentially followed by convergence, or (3) elevational conservatism, in which ancestral elevational ranges are retained. However, the relative contribution of these processes to generating patterns of elevational overlap and turnover is unknown.

**Location:** Tropical mountains of Central- and South-America.

**Time period:** The last 12 myr.

**Major taxa studied:** Birds.

**Methods:** We collate a dataset of 165 avian sister pairs containing estimates of phylogenetic age, geographical and regional elevational range overlap. We develop a framework based on continuous-time Markov models to infer the relative frequency of different historical pathways in explaining present day overlap and turnover of sympatric species along elevational gradients.

**Results:** We show that the turnover of closely related bird species across elevation can predominantly be explained by displacement of elevation ranges upon contact (81%) rather than elevational divergence in allopatry (19%). In contrast, overlap along elevation gradients is primarily (88%) explained by conservatism of elevational ranges rather than displacement followed by elevational expansion (12%).

**Main conclusions:** Bird communities across tropical elevation gradients are assembled through a mix of processes, including the sorting, displacement and conservatism of species elevation ranges. The dominant role of conservatism in explaining co-occurrence of species on mountain slopes rejects more complex scenarios requiring displacement followed by subsequent expansion. The ability of closely related species to coexist without elevational divergence provides a direct and thus faster pathway to sympatry and may help explain the exceptional species richness of tropical mountains.

## 1 Introduction

Explaining the combination and diversity of species that co-occur within ecological communities remains a major challenge. In part, this is because the patterns of spatial overlap observed at the present depend not only on current ecological interactions between species, but also on historical processes operating over much longer timescales and that are beyond the reach of direct observation or experimental manipulation (Weber *et al*., 2017). These historical processes include speciation, niche evolution, dispersal, and range expansions (Mittelbach & Schemske, 2015), with differences in the dynamics of these processes, and the ecological factors controlling them, underlying different theoretical models for how communities are assembled.

One of the main models for how communities assemble is based on the idea that new species arising in spatial isolation (i.e. allopatry) diverge in their ecological niche (including habitat), and only those that happen to diverge sufficiently to minimise competition are able to co-occur when they come back into secondary contact (Pfennig & Pfennig, 2010; Stuart & Losos, 2013). According to this model, community assembly largely involves the ‘sorting’ of pre-existing variation arising due to geographically variable selection pressures. An alternative to the ecological sorting model, proposes that niche differences between co-occurring species arise upon secondary contact, with competition between species driving divergent selection and a displacement of their ecological niches (Brown & Wilson, 1956). As with ecological sorting, this ‘ecological displacement’ model (termed ‘character displacement’ when considering heritable traits) assumes that niche similarity limits co-occurrence, but differs in its predictions of when and why niche differences evolve. A final possibility, is that species may reassemble into communities whilst retaining their ancestral niche. This ‘niche conservatism’ model (Cadena *et al*., 2011) is expected to predominate if ecological niche overlap is not limiting, either because of other constraints on co-occurrence (e.g. dispersal limitation) or because species have diverged along alternative niche dimensions (Pigot *et al*., 2018).

A classic system and spatial parallel for studying these historical processes concerns the distribution of species along tropical mountain slopes. Tropical mountains are renowned for their exceptional diversity. For instance, in the tropical Andes over 800 species of birds can occur on a single mountain slope (Walker *et al*., 2006). Since most speciation events involve the geographic isolation of populations on different mountains (i.e. allopatry)(Price, 2008; Cadena & Céspedes, 2020; Linck *et al*., 2020), the main problem is to understand how these species subsequently assemble on the same mountain slope (i.e. sympatry). In this system, sympatry may involve species co-occurring at the same elevation, what is often referred to as syntopy. Alternatively, species may occupy different elevation ranges. Indeed, many species have very narrow elevation distributions (e.g. a few hundred metres in vertical distance) and replace one another across elevational gradients (Diamond, 1973; Terborgh & Weske, 1975). Previous studies have variously provided evidence that conservatism (Cadena *et al*., 2011), sorting (Cadena, 2007), and displacement (Freeman, 2015; Freeman *et al*., 2022) of elevational ranges may individually be involved in shaping these patterns of co-occurrence and turnover along mountain slopes, but progress in disentangling their relative contributions has been limited.

One problem is that different assembly models can lead to the same present day pattern. In particular, according to the elevational conservatism (EC) model, species living on the same mountain slope have overlapping elevation ranges because they retain the adaptations to specific environmental conditions inherited from their ancestor (Cadena *et al*., 2011) (Figure 1d). However, while some studies have argued that elevational conservatism is a pervasive process (Linck *et al*., 2021), the same pattern could also arise under the elevational displacement (ED) model. Under ED, competition on secondary contact first forces species to occupy different elevations, but as species diverge along alternative niches dimensions (e.g., resource or microhabitat use) they may subsequently expand their elevation ranges and overlap along the gradient (Figure 1c). This ED model has long been regarded as the dominant process explaining the build-up of species within elevation zones on tropical mountains (Diamond, 1973; Freeman, 2015).

**Figure 1.**
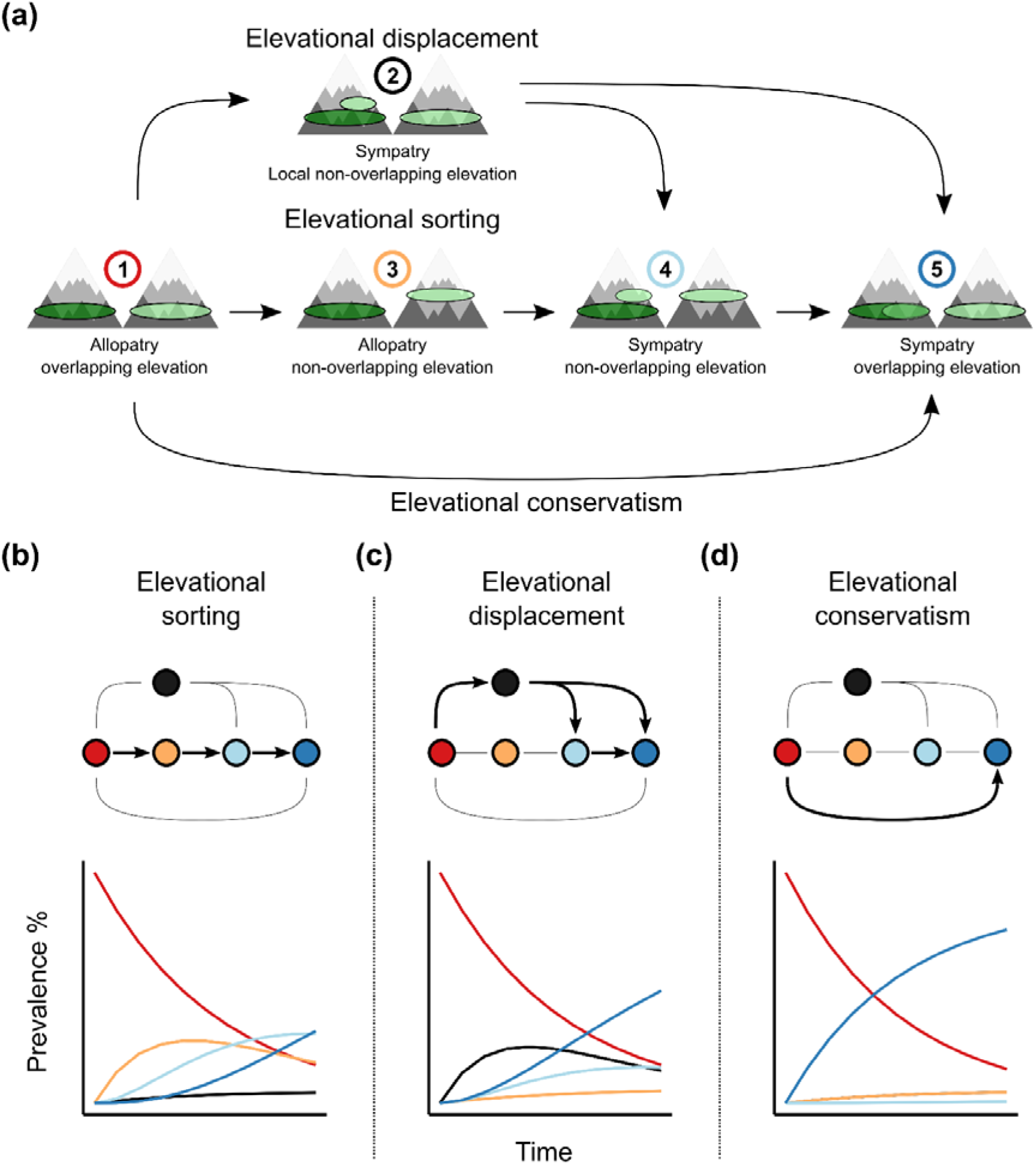
Hypothetical pathways towards sympatry in montane sister species. a) All possible transitions (arrows) in our model between the different states (numbers) are shown. States are defined by the geographical and elevational patterns of overlap between sister species ranges (coloured ovals) on mountains. Species pairs are assumed to start in a state of allopatry with overlapping elevational ranges (i.e. state 1). From this initial state, pairs may follow different possible community assembly pathways depending on which states they transition through: Elevational sorting (ES), Elevational displacement (ED) and Elevational conservatism (EC). b-d) The relative dominance of each pathway is expected to leave distinct signatures in how the prevalence of the five states across sister pairs changes with time since divergence. Here, for illustrating these distinct signals, curves of the expected prevalence are based on simulations in which 90% of sister pairs are assumed to pass through the (b) ES, (c) ED and (d) EC pathways. Note that while in (a) state 2 is represented as the shift in the elevation range of one sister species, both species could shift or contract relative to their range in allopatry.

A similar challenge exists when trying to discriminate elevational displacement from elevational sorting (ES). Both of these models predict that species living on the same mountain slope should have non-overlapping elevation ranges, at least prior to any subsequent expansion in elevation range. Thus, discriminating between these pathways depends critically on inferences about whether elevational divergence precedes (i.e. ES) or coincides with (i.e. ED) the attainment of secondary contact (Figure 1b, c). Studies attempting to address this issue have variously used phylogenetic information to compare the relative age of sister pairs that are elevationally divergent and those that are sympatric, or have compared elevation divergence between sister species in zones of sympatry to places where they remain allopatric (Freeman, 2015; Cadena & Céspedes, 2020; Figure S1). However, while such tests can provide evidence that elevational displacement does or does not occur, critically they do not reveal how frequently this process occurs and thus its relative contribution to the build-up of sympatric diversity and elevational turnover (Anderson & Weir, 2021). Progress in understanding community assembly, thus requires new approaches that can reliably infer the relative frequencies of different historical processes underlying current patterns of species distributions.

Here, we investigate the relative frequency of elevational displacement, sorting, and conservatism and how they contribute to the turnover and co-occurrence of passerine birds across elevation gradients in the Neotropics. We first compiled a new dataset consisting of the divergence time, geographical and regional elevational distributions of *n* = 165 sister species. We then apply a novel modelling approach that builds on previous studies examining the transition to sympatry using continuous-time Markov models (Pigot & Tobias, 2013; Pigot *et al*., 2018), but extend this framework by considering the multiple pathways through which species can attain sympatry via the evolution of elevational ranges. Focussing on avian sister species, where we can confidently assume that speciation involves the geographic isolation of populations (Phillimore *et al*., 2008; Price, 2008), allows us to use information on their current age and state (e.g. allopatric or sympatric, and overlapping in elevation or not) to model the historical pathways through which patterns of geographic and elevational range overlap arise. Using this approach, we address three main questions. First, what is the relative importance of elevational sorting, displacement and conservatism in the build-up of sympatry among Neotropical montane sister species? Second, what is the relative importance of elevational sorting and displacement in generating turnover between sister species across elevation gradients? Third, how does co-occurrence between species at the same elevation on the same mountain slope arise? We address these questions by modelling the historical pathways through which current distributions have arisen and conduct sensitivity tests and simulations to identify potential biases arising from limitations in the data or methods. Our results show that it is possible to reliably recover the historical pathways through which present day patterns of sympatry and co-occurrence have arisen, and we provide estimates of the relative frequency of these different pathways in explaining the assembly of Neotropical montane bird communities.

## 2 Material and methods

### 2.1 Elevational and geographic data

We compiled a dataset consisting of avian sister species occurring in the mountain ranges of the Neotropics. Sister species and their respective divergence times were extracted from two recently published phylogenies covering respectively >95% and >98% of the oscines and suboscines from North and South America (Barker *et al*., 2015; Harvey *et al*., 2020). We only retained sister pairs that met the following conditions: (1) At least one of the species occupies montane habitat, defined as areas >500m above sea-level (Freeman, 2015). (2) Both species occupy predominantly humid environments such that species pairs could realistically live on the same mountain slope and macrohabitat (e.g. we removed species pairs if one sister occupied predominantly arid habitats and not humid forest) as assessed with information from field guides. (3) Both species live in the Neotropics, which included the tropical mountain ranges of Central and South America. Species that had ranges stretching into North America were included in the dataset if they were primarily Neotropical. The final dataset consisted of *n* = 165 sister pairs.

We scored whether each sister pair was allopatric or sympatric and whether they had overlapping or non-overlapping elevational distributions using geographical and elevational range data gathered from a number of sources, including regional field guides and surveys (Hilty & Brown, 1986; Ridgely & Greenfield, 2001; Hilty, 2003; Walker *et al*., 2006; Vallely & Dyer, 2018). We supplemented this with *Birds of the World (BW)* (Billerman *et al*., 2020) that typically reflects information found in regional field guides but incorporates more recent taxonomic changes. We note that, although not primary scientific literature, field guides offer a wealth of expert knowledge on the natural history of birds, and are frequently used in studies of bird elevation distributions (e.g. Quintero & Jetz, 2018).

For each species, we used these sources to record elevational ranges on a region-by-region basis, usually corresponding to countries, but in greater detail where available (e.g. Choco vs. Amazonian slope in Colombia). This is important, not only to detect differences in elevation range overlap between zones of allopatry versus sympatry, but also because species can vary in their elevation limits across their geographic range more generally due to differences in climate, mountain height and biotic interactions (Janzen, 1967; Londoño *et al*., 2017; Pujolar *et al*., 2022). In some cases, elevation range limits in a particular region were not available, but field guides would report that elevational ranges of the sister species were non-overlapping. We used this expert information when assigning species as overlapping or not in that region. Observations of single specimens at single elevations were not incorporated in our dataset.

### 2.2 Assigning geographic and elevational range overlap

Although sister species can overlap in their geographic and elevational range to varying degrees, our modelling approach requires treating overlap as a discrete state. We defined species as having overlapping elevation ranges if overlap (range in vertical height, metres) was ≥20% of the species with the narrower elevation range. This threshold was chosen to avoid classing species which only meet marginally along narrow contact zones as overlapping. Sympatry was defined as when sister species were present on the same mountain slope, regardless of whether the geographic extent of overlap was widespread (i.e. thousands of kms) or minimal (i.e. a few kms). We did not use metrics of absolute or proportional geographic range overlap to define sympatry because this is unsuitable for montane systems where one species can have a very small geographic range. In practice, defining sympatry was unambiguous as allopatric pairs typically occurred on different mountain summits or regions separated by obvious geographical barriers (e.g. valleys).

We assigned species pairs to one of five possible discrete states, defined by the combination of sympatry/allopatry and elevation overlap/non-overlap (Figure 1a). For sister species occurring in allopatry, they may have overlapping (state 1) or non-overlapping (state 3) elevational ranges. Sister species occurring in sympatry may have non-overlapping elevational ranges (state 4), overlapping elevational ranges (state 5), or elevational ranges that overlap in allopatry but not sympatry (state 2). Pairs were not classified in a separate state if they showed elevational divergence in allopatry but not sympatry (opposite of state 2). Under this scenario, sympatry has been established irrespective of elevational differentiation, making the distinction of limited relevance to our analysis (and classified in state 5). Regardless, under the protocol discussed below merely one pair would be classified in such a state.

We found that for most species, variation among regions in their elevation range limits was relatively minor (Figure S2). However, we incorporated the regional variation that does exist, by using the mean pairwise overlap between sister species elevation ranges among the regions where they occur, doing this for the allopatric and sympatric parts of their geographic ranges separately. Thus, allopatric sister species are assigned to state 1 if their average regional elevation overlap is ≥20%. Otherwise, allopatric pairs are assigned to state 3. If pairs are sympatric, they are assigned to state 2, 4 or 5 according to the following rules. Sister species are assigned to state 2 if they show on average ≥20% overlap in elevation ranges between allopatric regions, and <20% overlap between sympatric regions. If sister species show on average <20% overlap between their respective allopatric ranges compared to the other species’ entire range, as well as an average overlap in sympatry of <20%, they are assigned to state 4. All remaining pairs, which show on average ≥20% elevation overlap in both sympatry and allopatry, are assigned to state 5. We refer to this method of using the mean pairwise regional overlap as the ‘average overlap’ protocol to distinguish this from alternative approaches used in our sensitivity analyses.

### 2.3 Sensitivity analyses

To ensure our results were robust to the way elevation range overlap was defined we performed a series of sensitivity tests. First, we repeated our analysis using 1 and 40% thresholds to define sister pairs as overlapping (Figure S3, S4, S5, S6). Second, we accounted for regional variation in elevation ranges in different ways. For zones of allopatry and sympatry we used: (i) the maximum upper and minimum lower elevational range limit of a species across regions to define its elevation range (ii) the minimum overlap observed among regional elevation ranges and (iii) the maximum overlap observed among regional elevation ranges. These different protocols which we refer to as the ‘global’, ‘minimum’ and ‘maximum’ approach, will variously tend to increase (‘global’ and ‘maximum’) or decrease (‘minimum’) the number of sister pairs assigned as having overlapping elevation ranges (Figure S3, S4, S5, S6). Third, we explored different ways of detecting the displacement of elevation ranges in sympatry (Figure S7).

In our main analysis we assigned sister species to state 2 if elevation ranges overlap in allopatry but not (i.e. <20%) in sympatry, a pattern expected if competition causes a displacement of species elevation ranges on secondary contact. However, this is a rather strict definition because displacement could be substantial but still not eliminate overlap. For example, sister species elevation ranges could overlap by 90% in allopatry and 50% in sympatry, a decrease in overlap indicative of competitive displacement but one that would not be detected by our assignment. To address this we perform an additional analysis in which species are assigned to state 2 if they show ≥20% reduction in overlap in sympatry compared to allopatry (Figure S7).

### 2.4 Statistical analysis using Markov models

To infer the relative importance of different community assembly pathways in explaining patterns of geographic and elevation overlap and turnover, we constructed a continuous time multi-state Markov model (Figure 1b). In this model, the initial state for sister species is allopatry with overlapping elevational ranges (state 1), which reflects the situation expected at the time of their initial divergence (Coyne & Orr, 2004). Sister pairs then stochastically transition between states, with different transitions corresponding to the different assembly pathways (Figure 1). Under the EC pathway, sister species transition directly from state 1 to state 5 (i.e. sympatry and overlapping elevational ranges; Figure 1d) while under the ED and ES pathways species must pass through intermediate states. Under the ED pathway, sister species transition from state 1 to state 2 (i.e. retaining overlapping elevation ranges in allopatry but non-overlapping elevation ranges in sympatry), before then converging or expanding in elevation in sympatry to reach state 5 (Figure 1c). Under the ES pathway, sister species transition to state 3 (i.e. non-overlapping elevation ranges whilst remaining in allopatry), then state 4 (i.e. attaining sympatry and retaining non-overlapping elevation ranges) and finally to state 5 (Figure 1b). Thus, our model allows for the elevation overlap of sister pairs in sympatry (state 5), which is the final absorbing state, to be attained through any of the three community assembly pathways.

Within this framework, we considered two additional scenarios. First, in our model we also allow sister pairs in state 2 to transition to state 4, representing a process where species that have attained sympatry via elevation range displacement subsequently diverge in their elevation range in the allopatric part of their distribution (Figure 1c). Such divergence in allopatry, was hypothesised by Diamond (1973) to be the result of the spread of adaptations to the compressed elevational range arising in sympatry, resulting in displaced elevation ranges even where the sister species is not present. Second, in an additional analysis we relax the assumption that all sister pairs arise in state 1, by estimating an additional parameter, *γ*, representing the probability that species arise in state 3 (Supplementary Information Figure S8, S9). This accounts for the possibility that newly formed sister species may immediately occur at different elevations (e.g. due to differences in climate and thus heights of habitat zones between regions).

Based on the estimated time since divergence and states of sister pairs at present we used maximum likelihood (ML) (Jackson, 2011) to estimate the transition rates between the states. The full model contains 7 rate parameters corresponding to the 7 possible state transitions. We also considered simpler models by constraining combinations of parameters to have identical rates. The simplest model has only a single rate parameter, corresponding to identical rates for all transitions. We used AIC to compare model fit across all (*n* = 877) possible model simplifications and report both the best model and the model-averaged parameter values of all highly supported models (ΔAIC ≤ 2 of best model).

### 2.5 Relative frequency of community assembly pathways

Having inferred the transition rates between states, we then used these to estimate the relative contribution of elevational displacement (ED), elevational sorting (ES), and elevational conservatism (EC) to pairs leaving state 1 and arriving in states 2, 4 and 5. Specifically, we used the inferred rates to perform 1000 posterior predictive simulations using the Gillespie algorithm for constant rates (Gillespie, 1977). Under the Gillespie algorithm, transitions between states correspond to events. The simulation starts at time *t* = 0, indicating the time at divergence for all sister species. The waiting time (*δ*) to the next event is determined by a random draw from an exponential distribution with the mean equal to the sum of all transition rates across all sister pairs. For example, if the estimated transition rate from state 1 to 2 (r12) is 0.05 and there are currently 5 pairs in state 1, then this transition adds 0.25 to the total rate. Transition rates are assumed to be constant through time, but older species will have more time to experience a transition. The event that occurs at time *t + δ* is selected with a probability equal to the relative contribution of each rate to the total rate. This transition applies to a single species pair and this pair is chosen at random with equal weighting across all pairs that are currently in the relevant state. As we simulate forward in time, species can no longer transition if their age is less than *t* and thus the simulation continues for a period equal to the age of the oldest sister pair. During the simulation we record the percentage of sister pairs passing through each of the three different community assembly pathways (i.e. ED, ES, and EC), and report the median and 95% confidence intervals (CI) in these %’s across simulation runs.

### 2.6 Assessing model fit

Although the model is optimised using maximum likelihood, model fit may be poor if the underlying assumptions of the model are not met. If that is the case, estimated rates will poorly reflect the empirical observations. We assessed how well the predicted transition rates can predict (*i*) the change in frequency of states through time, and (ii) the distribution of sister species divergence times for each state using the output of the posterior predictive simulations. To examine if the model can adequately predict state changes through time we binned species pairs in three equal sized age bins of 55-56 sister pairs each. We used three bins to ensure we would approximately capture nonlinear changes in the prevalence of states through time beyond just increases and decreases. The prevalence of each of the five states per bin is then compared between the empirical data and 95%CI’s constructed from the final prevalence of the simulated states. If the empirical prevalence falls within the 95%CI this would indicate good model fit (Figure 3a), but large CI’s likewise indicate high uncertainty. Finally, we also compared the empirical age distribution of each state to the average distribution of ages across the posterior simulations (Figure 3b).

### 2.7 Simulation tests of accuracy and precision

Using simulations, we further evaluated the model by assessing if we can both accurately and precisely recover the true transition rates. High accuracy indicates that the model is not biased towards over-or under-estimating transition rates. High precision indicates that the estimated rates are close to the true rates. We explored multiple scenarios (Supplementary Information Table S1-S4), designed to characterise the three different community assembly pathways (S1-S3), as well as a scenario assuming identical transition rates among states (S4) and one corresponding to the transition rates inferred from the model-averaged fit to the empirical data (S5). For each scenario, we performed 100 replicate simulations using the observed number and ages of sister pairs. For each simulation, we then performed an identical model fitting procedure as for our empirical data, resulting in transition rate estimates according to the best and model-averaged approach for each simulation. We constructed 95% CIs from the estimates across the simulations for the best and model-averaged approaches to evaluate the accuracy and precision of the rate estimation. Additionally, we determined the coverage of the models (*sensu* Morris *et al*., 2019) by inferring how often the (a) 95%CI of the best model and (b) unconditional CI (Anderson, 2008) of the model-averaged estimates for an individual simulation captured the true rates as predetermined for every scenario (‘rate coverage’). We also investigated how potential error in rate estimates impacts the estimation of the relative frequency of community assembly pathways taken by sister pairs (‘pathway coverage’, Table S3, S4).

## 3 Results

### 3.1 Empirical distribution of sister pair states

Across the *n* = 165 sister species pairs, the majority are currently in allopatry with overlapping elevation ranges (state 1: 58%). The next most common state is sympatry with overlapping elevation ranges (state 5: 18%), with fewer pairs having non-overlapping elevational ranges and occurring in sympatry (state 4: 7%), or allopatry (state 3: 8%), or having non-overlapping elevational ranges in sympatry but overlapping elevational ranges in allopatry (state 2: 10%). The median age of pairs that are in allopatry with overlapping elevation ranges (state 1: 2.09Myr) is younger than all other states (state 2: 3.36, state 3: 2.74, state 4: 3.98, and state 5: 2.73Myr), consistent with our assumption that it is the initial state at the time of species divergence.

### 3.2 Transition rates between states

We found that our best model contains two parameters and constrains r12, r13, r15, r24, r25, and r45 to 0.07 (95% CI: 0.06-0.09), and r34 to 0.28 (95% CI: 0.13-0.63) transitions per pair per million years (Myr^-1^). While there are 66 models that are highly supported (≤2 ΔAIC), the model-averaged rate estimates are very similar to the best model. Because of this, we focus on the model-averaged results below. The model averaged results show that the transition rate from the initial state of allopatry with overlapping elevation ranges (state 1) to non-overlapping elevation ranges, either while in allopatry (r13 = 0.06 Myr^-1^) or upon the attainment of sympatry (r12 = 0.08 Myr^-1^) is relatively slow, and similar to attaining sympatry while conserving elevational ranges (r15 = 0.08 Myr^-1^). Once elevational differentiation has occurred in either allopatry (state 3) or upon secondary contact (state 2), the transition to elevational differentiation in both allopatry and sympatry (state 4) is relatively fast (r34 = 0.18 Myr^-1^ and r24 = 0.17 Myr^-1^). In contrast, the transition from a state where species have non-overlapping ranges in sympatry (state 2 and 4) to one where they co-occur at the same elevation in sympatry (state 5) is relatively slow (r25 = 0.06 Myr^-1^ and r45 = 0.08 Myr^-1^).

### 3.3 The relative contribution of community assembly pathways

The posterior-predictive simulations show that following speciation a similar proportion of pairs embark on the ES (28.36%, 95% CI: 18.52-38.89%), ED (35.06%, 95% CI: 24.66-47.30%) and EC pathways (36.00%, 95% CI: 25.71-45.95%) (Figure 2b). Of the pairs with currently non-overlapping elevation ranges in sympatry (states 2 and 4), substantially more are inferred to be generated through ED (81.48%, 95% CI: 65.51-95.83%) than ES (18.52%, 95% CI: 4.17-34.49%) (Figure 2c). Of the species pairs that attain overlapping elevation ranges in sympatry (state 5) almost all of these (88.46%, 95% CI: 74.98-97.68%) attain this state via the EC pathway rather than the ES (0.00%, CI: 0.00-9.09%)or ED (9.09%, 95% CI: 0.00-21.74%) pathways (Figure 2d). This can be explained because although a similar proportion of pairs embark on each pathway (i.e. r12 ≈ r13 ≈ r15), species pairs taking the ES (1⍰3⍰4⍰5) and ED (1⍰2⍰4⍰5 or 1⍰2⍰5) pathway must pass through a number of intermediate states to attain overlapping elevation ranges in sympatry and this takes much longer than the direct EC pathway (1⍰5). We note that these key results remained qualitatively the same when using (i) alternative thresholds for defining elevation range overlap, (ii) different approaches for incorporating regional variation in elevation ranges, (iii) different criteria for detecting elevational displacement on secondary contact and (iv) relaxing assumptions about the initial state of sister pairs at speciation (Figure S3, S4, S5, S6, S7, S8).

**Figure 2.**
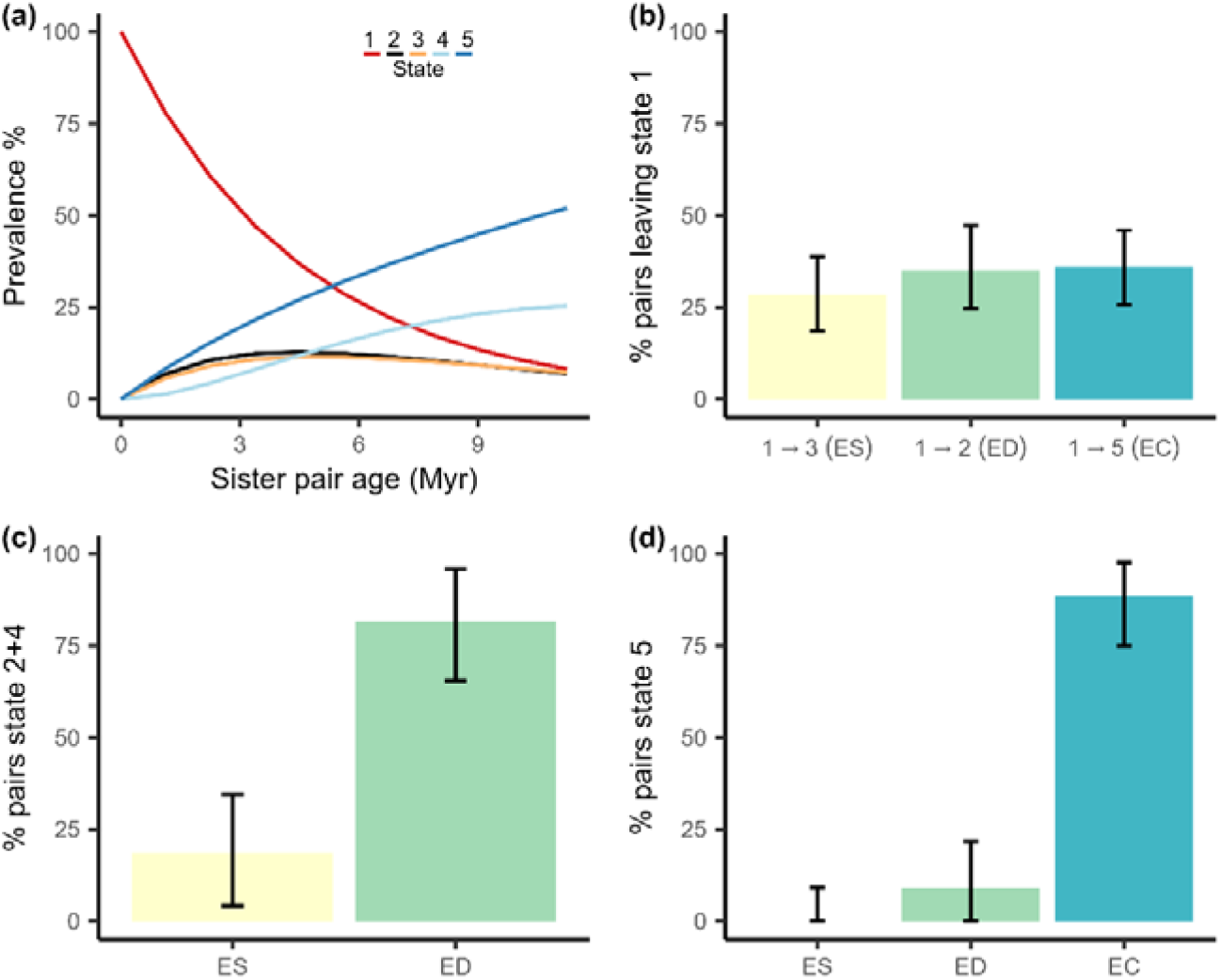
The inferred contribution of each community assembly pathway in explaining patterns of sympatry, turnover and overlap among sister species. a) estimated prevalence of the five states with time since divergence (Myr), as represented in Figure 1 but based on rates inferred by our model from the empirical data. Note the prevalence of state 2 (black) and 3 (yellow) are very similar. b) Percentage of sister pairs that have left state 1 and have transitioned to state 3, 2, or 5 representing the elevational sorting (ES), displacement (ED), or conservatism (EC) pathways respectively. c) the percentage of pairs that currently occupy non-overlapping elevational ranges in sympatry (states 2 and 4) that have arisen through the ES or ED pathway. d) The percentage of pairs that are sympatric with overlapping elevational distributions (state 5) that have arisen through the ES, ED, or EC pathway. Results are obtained from posterior-predictive simulations of the model-averaged parameter estimates and using a threshold of 20% overlap and the ‘average overlap’ protocol to define elevational overlap.

### 3.4 Assessing model fit and the accuracy and reliability of estimated transition rates

Our simulations show that transition rates from state 1 (r12, r13, and r15) (Figure 1), can be reliably and accurately estimated irrespective of the simulation scenario (Table S1, S2). Accordingly, our simulations also show that we can reliably recover the contribution of ED and ES pathways in generating sympatric sister pairs with non-overlapping elevation ranges (states 2 and 4) (Table S3-S4). Later transitions (r24, r25, r34, r45) are estimated with less accuracy, probably because there are relatively few old sister pairs and thus less information to reliably estimate these rates. As a result, when it comes to explaining the species that are currently sympatric and have overlapping elevation ranges (state 5) there is a bias towards overestimating the contribution of ED and underestimating EC (Table S3, S4) (Figure 2d). We note that this does not influence our conclusions, because despite a bias against detecting EC, we still infer this as by far the most common pathway to state 5. Overall, our simulations show that we can predict the incidence of the five states well through time (Figure 3). The simulations capture the main patterns, namely: an increase with age of sympatric pairs with overlapping elevation ranges (state 5), a decrease in allopatric pairs with overlapping elevation ranges (state 1), and slight increases for pairs that are either in allopatry or sympatry but have diverged in elevation (state 2, 3 and 4).

**Figure 3.**
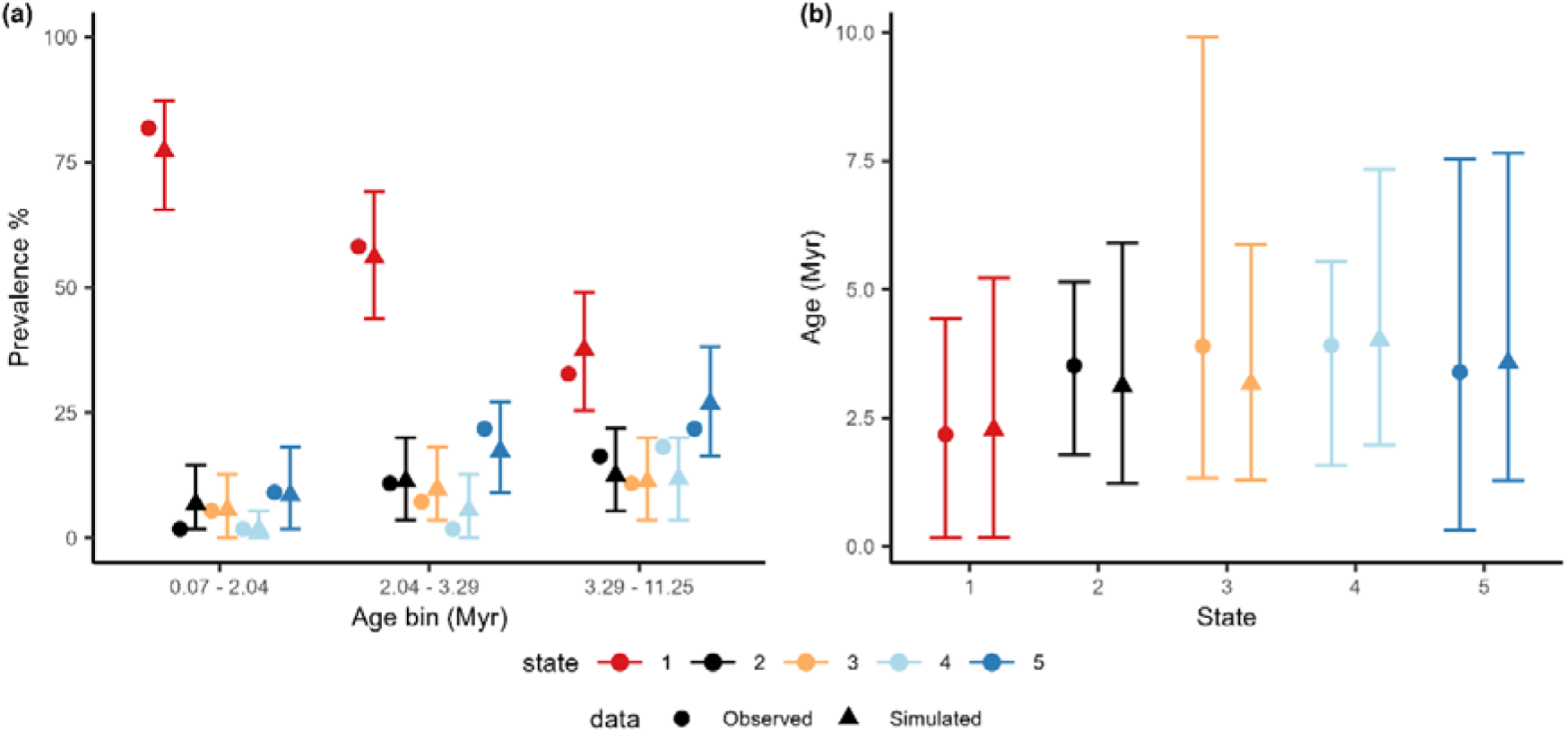
Tests of model adequacy. Empirical and predicted prevalence of states through time and age distributions of states. a) Empirical (circles) and predicted (triangles) prevalence (triangles) represent the prevalence of the states within three age bins of approximately equal size (n = 56, 55, and 55). Brackets represent the 95%CI of the prevalence of each state across 1000 posterior-predictive simulations. b) Empirical and predicted age distributions per state. Brackets indicate the mean and 95%CI for the empirical age distributions and the average mean and 95%CI over 1000 simulations for the predictions. Results are obtained from the model-averaged parameter estimates and using a threshold of 20% overlap and the ‘average overlap’ protocol to define elevational overlap.

## 4 Discussion

Multiple historical processes have been proposed to explain the patterns of range overlap and turnover across tropical elevational gradients. However, because previous studies have focused on static biogeographic patterns and treated elevational sorting, displacement and conservatism as alternative explanations, their relative contribution has remained unknown. Through our analysis of the dynamics of sympatry and elevation overlap among Neotropical montane birds, we present three key findings. First, we show that the relative contributions of different historical assembly models can be reliably inferred given current phylogenetic and geographic data among sister species. Second, our results show that elevational displacement upon secondary contact (>81%) contributes substantially more to the turnover of closely related bird species across elevation gradients than sorting (<19%) of species that diverged to occupy different elevations in allopatry. Finally, we show that the overlap of sister species along elevation gradients is almost entirely (>88%) explained by species attaining sympatry while retaining their ancestral elevational range, rejecting more complex scenarios requiring displacement upon secondary contact followed by subsequent shifts to occupy the same elevational range.

The high biodiversity of tropical mountains is associated with rapid turnover in community composition, as ecologically similar species replace one another up the mountain. Previous studies of montane birds have argued that elevational displacement rather than sorting would be the dominant process explaining such elevational replacements, but these studies have been unable to quantify the relative contribution of these pathways (Diamond, 1973; Freeman, 2015; Freeman *et al*., 2022). Our results show that both sorting and displacement contribute to the turnover in sister species across elevations, and that current patterns of sympatry cannot be explained by either process acting alone. However, we also show that elevational displacement upon contact is far more common that sorting, accounting for ∼81% of sympatric sister pairs with non-overlapping elevation ranges. These results are consistent with the idea that negative interactions between species (e.g. competition or apparent competition) cause displacement to occupy different elevations, and that this is the dominant process driving the layering of species across mountain slopes.

While our analysis shows that both elevational sorting and displacement are involved in generating turnover of sister species across elevation gradients, we find that sympatry is frequently reached without differentiation in elevation. We estimate that ∼45% of sister species living on the same mountain slope and ∼88% of species additionally living at the same elevation attain sympatry while conserving their ancestral elevation range. Such a high frequency of elevational range conservatism may not seem surprising given that previous studies focussed on the drivers of speciation have shown that most vertebrate sister species have overlapping elevation ranges (Cadena *et al*., 2011). However, a high frequency of elevation overlap among sister species is by itself inconclusive regarding how communities assemble because the same pattern can arise under the elevational displacement pathway if species subsequently expand their elevation ranges as they diverge across alternative niche axis (e.g. resource use) and competition is relaxed. Indeed, based on his studies of New Guinea birds, elevation displacement followed by expansion was the pathway proposed by Diamond (1973) for how diverse communities at a given elevation had assembled. Our phylogenetic approach to modelling the dynamics of elevation range overlap, enables us to exclude this possibility. Indeed, we estimate that displacement and subsequent overlap in species elevational ranges contributes little (9.09%) to current patterns of co-occurrence among sister species along elevation slopes.

The finding that a high proportion of allopatric sister species directly transition to occupy overlapping elevations in sympatry need not suggest that competition or other negative species interactions are unimportant in limiting coexistence for these species. Indeed, evidence that interspecific competition limits elevational ranges is widespread in birds (Terborgh & Weske, 1975; Freeman *et al*., 2022). Instead, species attaining sympatry without diverging in their elevation range may have diverged across alternative niche dimensions (either in allopatry or upon secondary contact) such as resource or microhabitat use. Such an explanation would be consistent with previous evidence that sympatry among Neotropical bird species is limited by divergence in key trophic traits, such as beak size (Pigot & Tobias, 2013; Pigot *et al*., 2018). Furthermore, while our analysis is limited to sister species and does not consider how patterns of elevation overlap and turnover arise among more distantly related lineages, it is among recently diverged species that competition is expected to be strongest (Pigot & Tobias, 2013). Therefore, while extending our models to deeper phylogenetic scales is an important direction for future research we do not expect this to overturn our finding that conservatism in elevation ranges is a more common route to co-occurrence than elevational range displacement followed by expansion.

In addition to the effects of phylogenetic scale, it is important to consider the possibility that the relative mix of different assembly processes may not be static over geological time. Given the relatively young age of many Neotropical montane radiations, and that there is little evidence for a slowdown in the rates of diversification (Weir, 2006; Harvey *et al*., 2020), local niche space at any point along the elevational gradient may currently be far from saturated. As niches become increasingly densely packed over time, it is possible that elevational sorting and displacement may become increasingly important pathways to sympatry as has been suggested for New Guinean (Diamond, 1973) and Himalayan birds (Price *et al*., 2014). It would therefore be interesting to apply our modelling approach to other organisms and tropical mountain systems to test whether the relative mix of different assembly pathways is consistent or varies across different contexts and stages of evolutionary radiations.

Fitting our model requires reliable information on both geographic and elevation overlap and for these spatial properties to each be encoded as simple binary states. In reality, elevation ranges can vary in complex ways, for example, showing overlap in some places but not in others, and with species shifting their elevation ranges across regions to match changes in climate and habitats. Here, we focussed on Neotropical birds occupying humid forests for which distributions are relatively well known and where we could incorporate regional variation in species elevation limits into our metrics of overlap. While we found that estimates of different community assembly pathways were robust to the way elevation range overlap was scored, accounting for the full complexity of overlap patterns may best be addressed using a spatially explicit model (Reijenga *et al*., 2021; Hagen, 2022). Such a model could incorporate the actual environmental conditions (and their variation across space) that limit geographic ranges as well as biotic interactions beyond those occurring between only sister species (Rangel *et al*., 2018; Hagen *et al*., 2021). Due to their complexity, such spatially explicit models are difficult to fit to empirical data, but the parameters that we estimate here could be used as priors to inform the rates of key processes (e.g. transition times to sympatry or elevation divergence).

In our model, sympatric species that co-occur at the same elevation (state 5) represent the final stage in the pathway of community assembly. The fact that not all sister pairs exist in this state is explained by new speciation events re-setting the cycle, so that insufficient time has passed since speciation for most sister pairs to transition to this state. Our model uses the observed distribution of sister pair ages which are determined by rates of speciation and extinction (Weir & Schluter, 2007), but it does not consider how the geographic or elevational overlap of species could in turn feedback to influence these diversification dynamics. Because speciation and extinction events prune or replace the set of sister pairs in a phylogeny, if these rates depend on the geographic or elevation overlap of a species, this could bias the estimated transition rates. For example, we speculate that sympatric species may have a higher instantaneous probability of speciating because the expansion of their geographic range makes them susceptible to renewed rounds of geographic isolation (Weir & Price, 2011). In this case, our model may underestimate transition rates to sympatry. While we think it is unlikely that such biases would also apply to elevation overlap, developing models that incorporate the reciprocal evolution between geographic/elevation overlap and diversification dynamics is an important avenue for future research (Goldberg *et al*., 2011; van Els *et al*., 2021; Landis *et al*., 2022).

The enormous diversity of species in tropical mountain ranges appear as stark outliers in statistical models seeking to explain global variation in species richness on the basis of climate and topography (Rahbek *et al*., 2019b). This is often attributed to the rapid turnover of species across elevations and mountains (Rahbek *et al*., 2019a). Our results provide a different perspective, showing that the rate of range expansions leading to sympatry are substantially accelerated by the capacity for species to occur on the same slope without having to first diverge in their elevation range. Specifically, the estimated mean lag time to sympatry following speciation is 6.01Myr, substantially shorter than the 9.46Myr that would be expected if the direct transition from allopatry to co-occurrence in sympatry at the same elevation (i.e. 1⍰5) were precluded (see Supplementary Information). Thus, in addition to the turnover of species across elevations or mountains, a key additional ingredient explaining the high diversity of tropical mountain ranges is the capacity for species to coexist locally without having to diverge into different elevational zones.

Our model represents a simplification of the complex processes governing the assembly of montane biotas and is limited to explaining the patterns of sympatry and elevation overlap among sister species. However, to our knowledge this is the first study to quantify the relative frequency of elevational sorting, displacement, and conservatism in shaping species distributions across montane gradients. A key next step will be to test how well our results generalise to different mountain regions or taxa that vary in their ecology. Our model could also be applied to disentangle the dynamics of assembly across other ecological gradients, such as the vertical layering of foraging niches among rainforest birds (MacArthur, 1958), perch height among Anolis lizards (Lister, 1976), or depth zonation in the Cichlid fish of East African rift lakes (Rodríguez & Lewis, 1997).

## Data Accessibility Statement

Data used in this study was compiled from previously published sources. Code and the data set are made publicly available on acceptance.

## Author contributions

Conceptualisation, XX1, XX3, XX4; data curation XX1, XX2; formal analysis, XX1; funding acquisition, XX4; investigation, XX1; visualisation, XX1; writing – original draft, XX1; writing – review and editing, XX1, XX2, XX3, XX4. All authors have read and agreed to submit the current version of the manuscript.

