## Supplemental figures and methods for "Disentangling the historical routes to community assembly in the global epicentre of biodiversity"

**Supplementary Material**

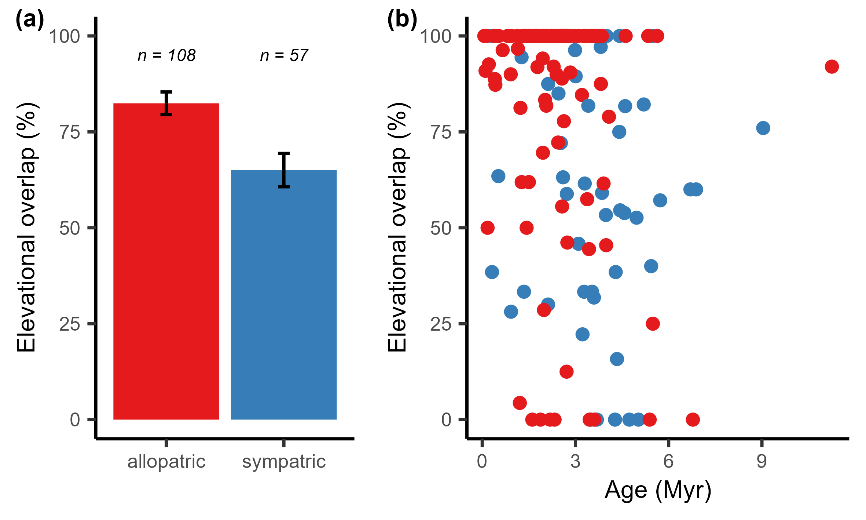

**Figure S1** Range-wide elevational overlap as a function of (a) geographic overlap and (b) time since speciation (Myr) across sister species pairs. Elevational overlap (%) is calculated using the maximum upper and minimum lower elevational range limit of a species across all regions (i.e. countries in the Neotropics) where it occurs (i.e. ‘Global overlap’ approach) and is shown for sister pairs that are allopatric (n = 108, red) and those that are sympatric (n = 57, blue). Brackets in (a) indicate the standard error.

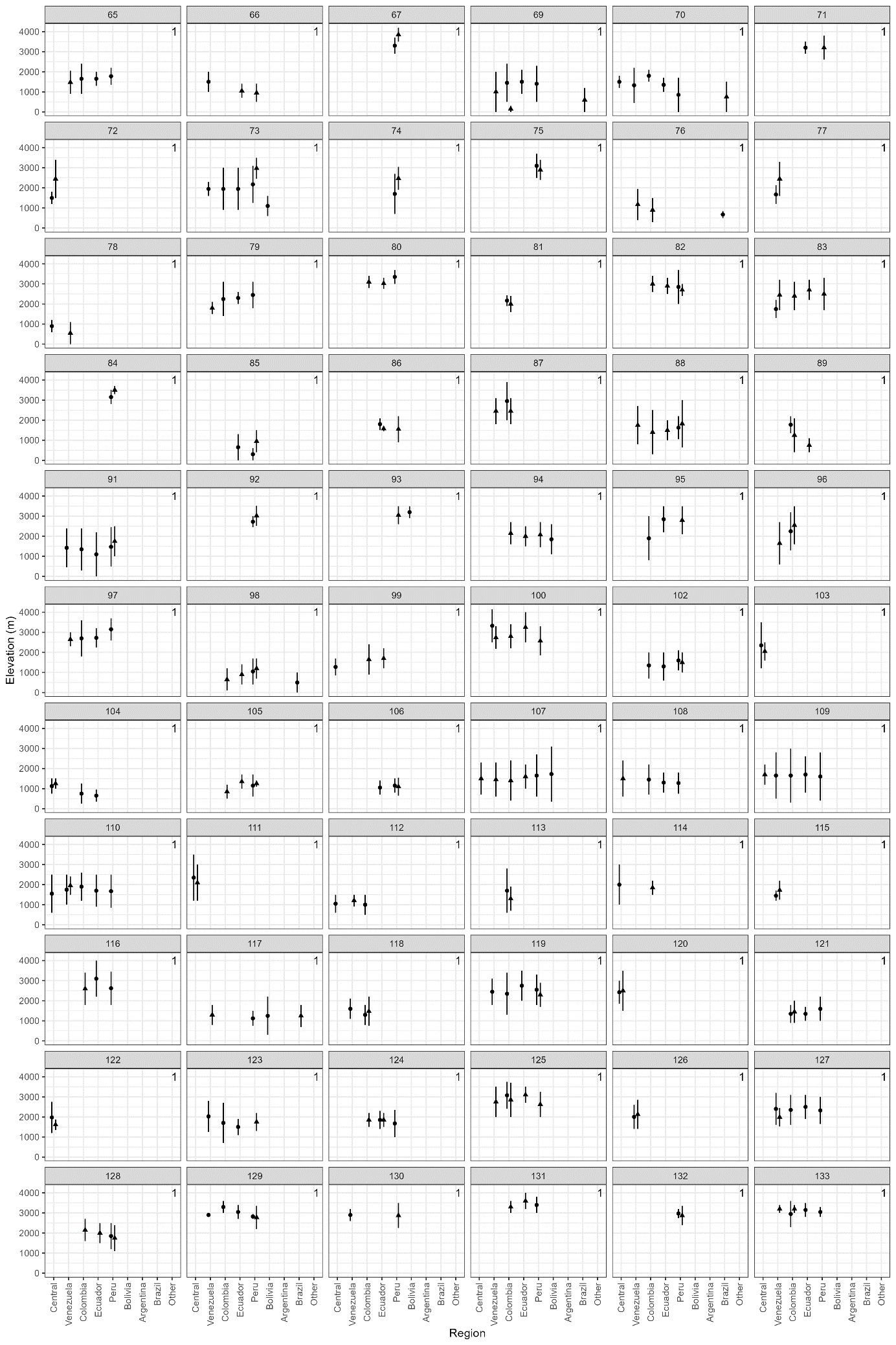

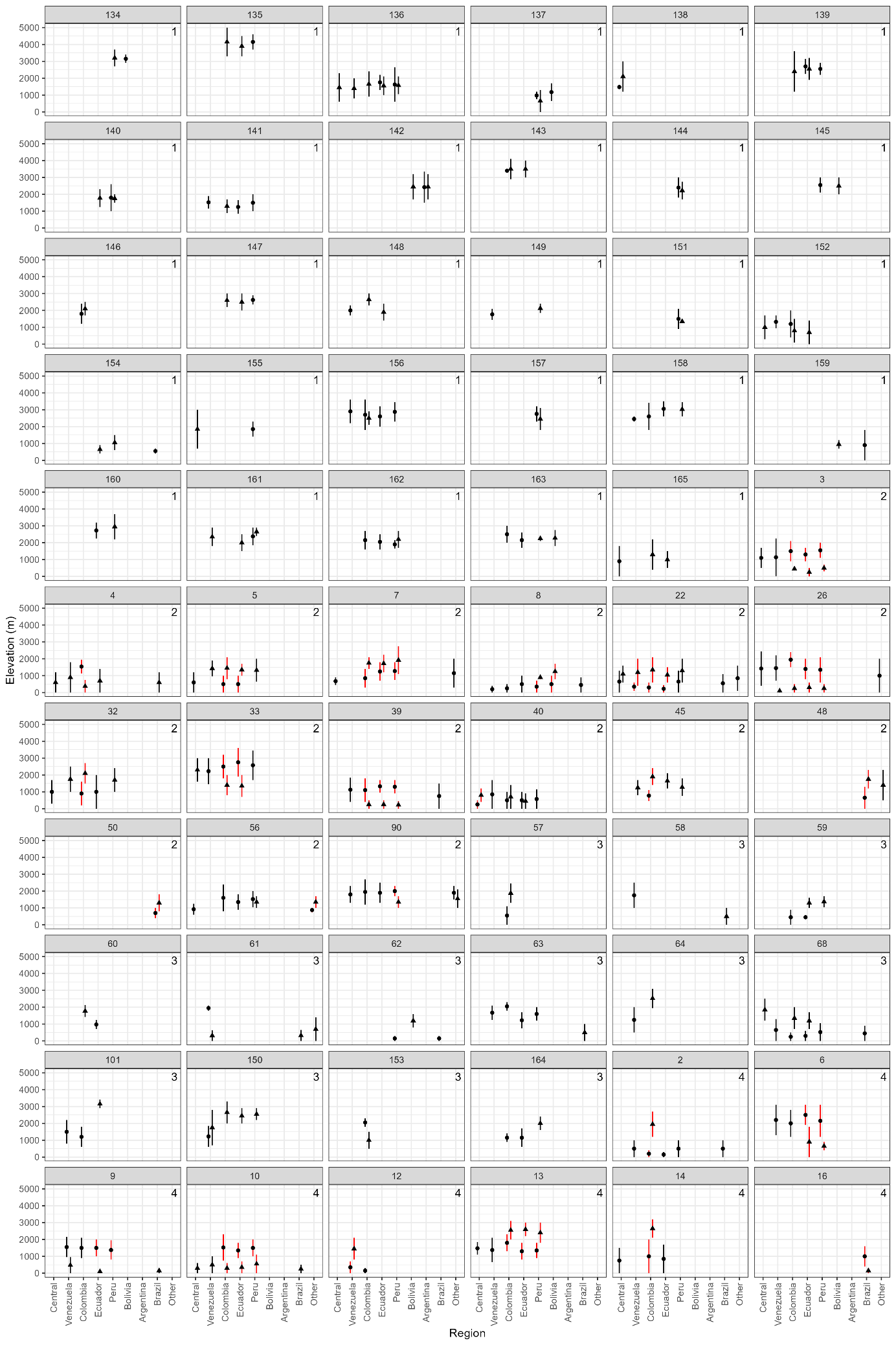

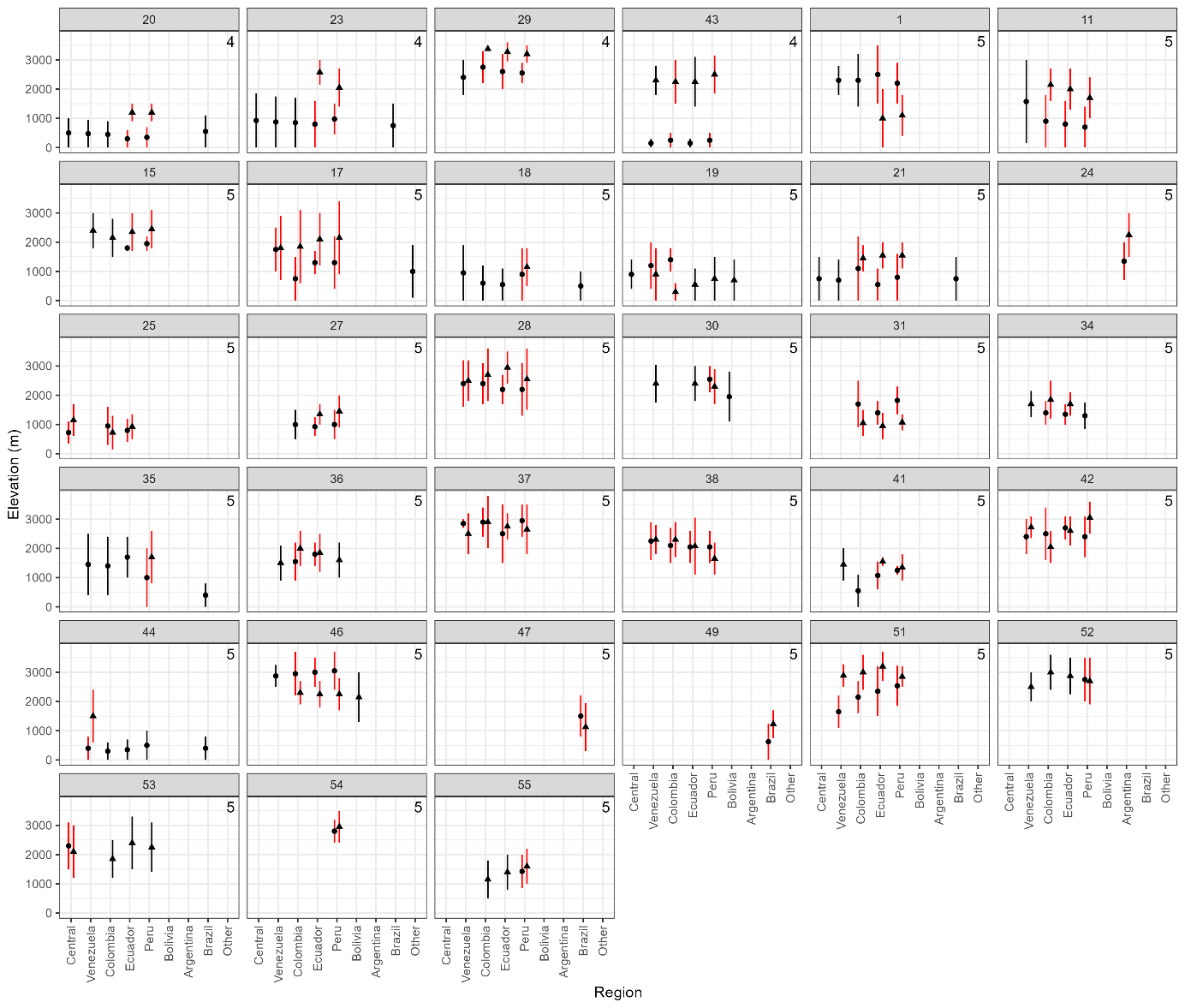

**Figure S2** elevational range limits per region for all sister pairs. Each panel shows one sister pair, with country level regions on the x-axis. “Other” refers to specific localities within regions for which information on elevational range limits was available. Bars show the highest and lower range limits of a species per region, with shapes indicating each sister species. States as described in Figure 1 are assigned based on the ‘average overlap’ protocol and shown in the upper-right corner of each plot. For species partially in sympatry (state 2, 4 and 5) red shows regions of sympatry. Note that both species can be present in a region without being sympatric. Numbers at the top of each plot are the unique identifier for each sister pair and correspond to those in Table S5 included in the supplementary data.

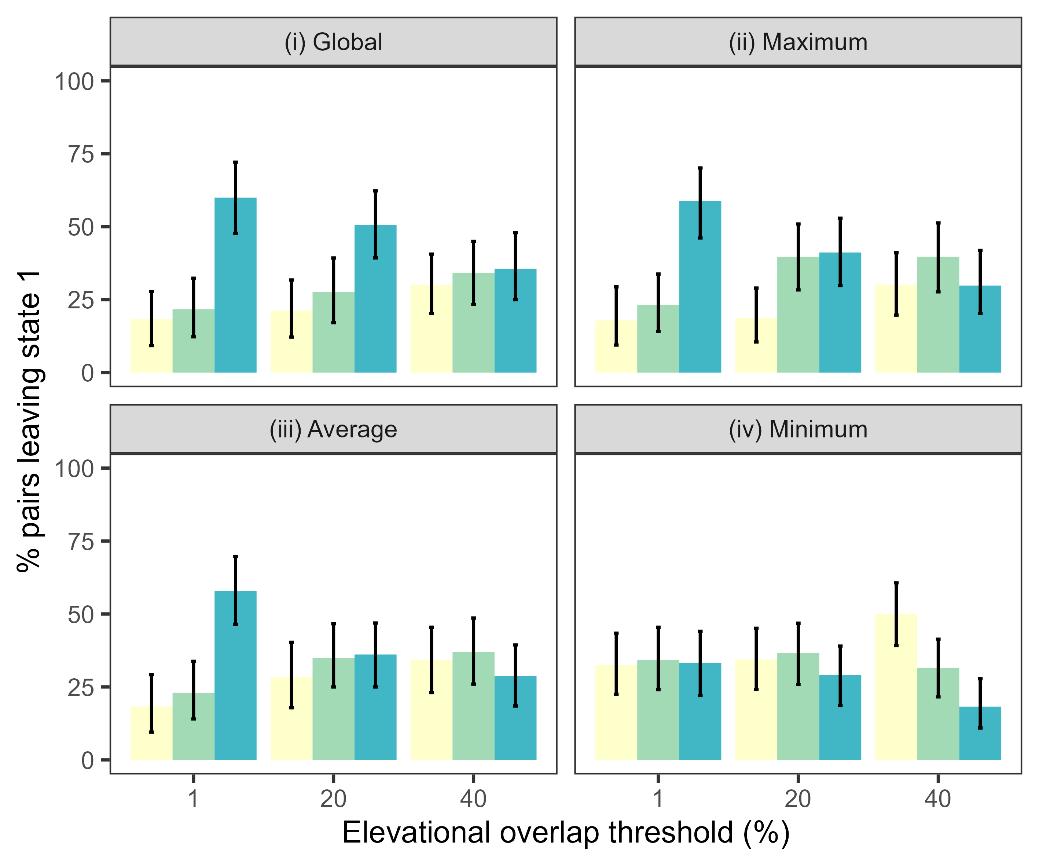

**Figure S3** Frequency of inferred community assembly pathways initiated in relation to the protocol used to determine elevation overlap (panels) and the % overlap threshold (1, 20 & 40%). Colours are as in Figure 2b (yellow; ES, green; ED, and blue; EC). The protocols respectively use (i) the global overlap based on the maximum upper and lower elevational range limits across all regions, (ii) the maximum pairwise overlap observed between regional elevational ranges, (iii) the average (mean) pairwise overlap, and (iv) the minimum pairwise overlap. The results in the main text are for the ‘average overlap’ protocol using a 20% threshold.

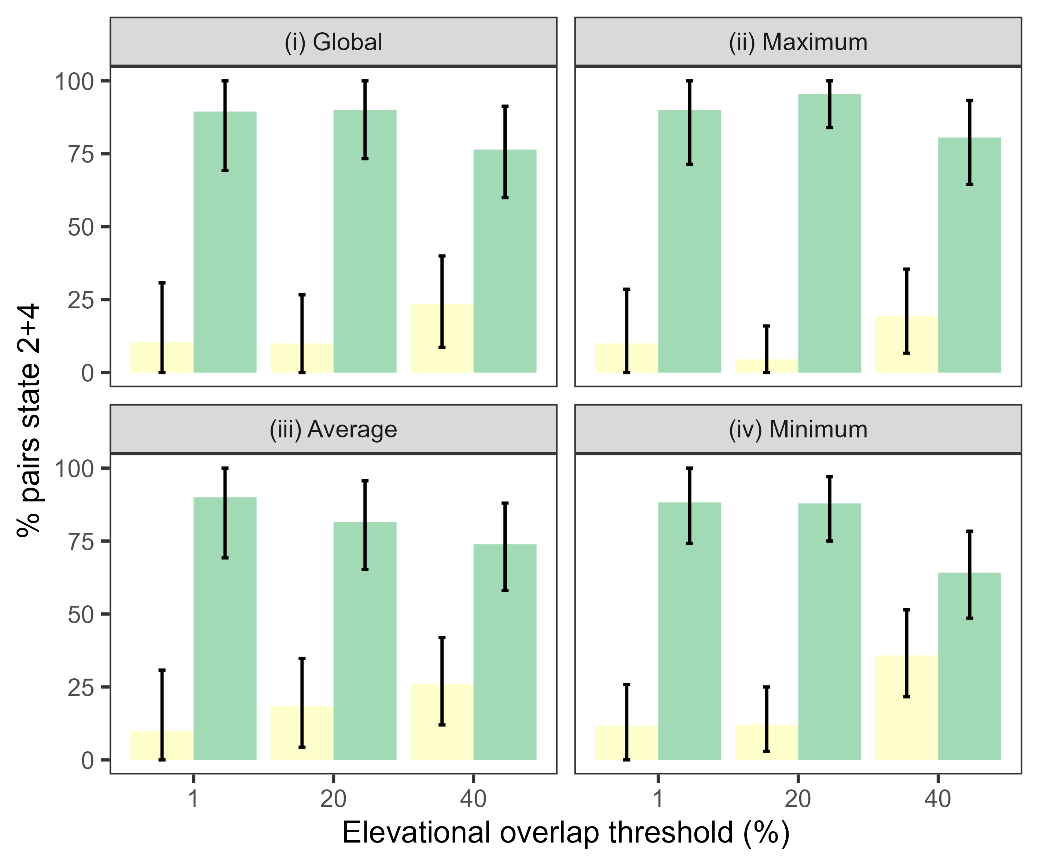

**Figure S4** Frequency of inferred community assembly pathways taken to explain elevational turnover (i.e. state 2 and 4) in relation to the elevational overlap threshold (1, 20 & 40%) and protocol (panels). Colours are as in Figure 2c (yellow; ES, green; ED). The x-axis shows the elevational thresholds, and each facet shows a protocol. The protocols respectively use (i) the global overlap based on the maximum upper and lower elevational range limits across all regions, (ii) the maximum pairwise overlap observed between regional elevational ranges, (iii) the (mean) average pairwise overlap, and (iv) the minimum pairwise overlap. The results as in the main text are as in the ‘average overlap’ protocol using the 20% overlap threshold.

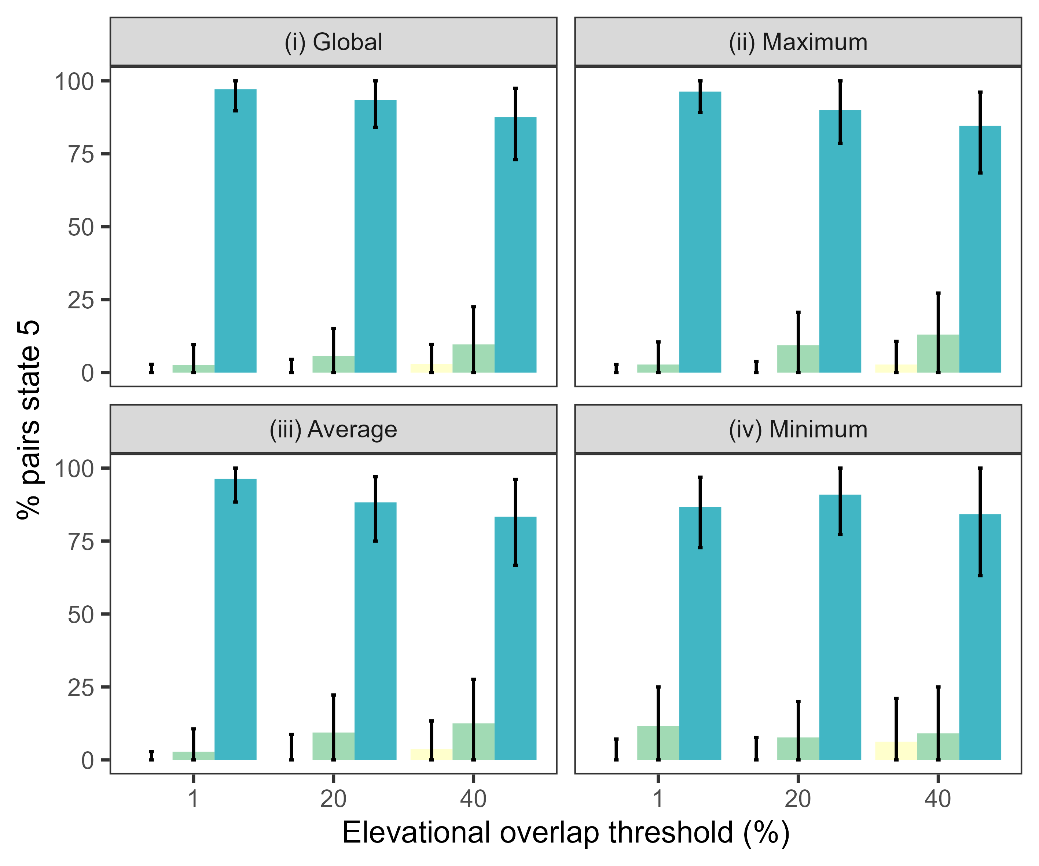

**Figure S5** Frequency of inferred pathways taken to sympatry with overlapping elevational distributions (state 5) in relation to the elevational overlap threshold (1, 20 and 40%) and protocol (panel). Colours are as in Figure 2d (yellow; ES, green; ED), and blue; EC). The protocols respectively use (i) the global overlap based on the maximum upper and lower elevational range limits across all regions, (ii) the maximum pairwise overlap observed between regional elevational ranges, (iii) the (mean) average pairwise overlap, and (iv) the minimum pairwise overlap. The results as in the main text are represented by the ‘average overlap’ protocol using the 20% overlap threshold.

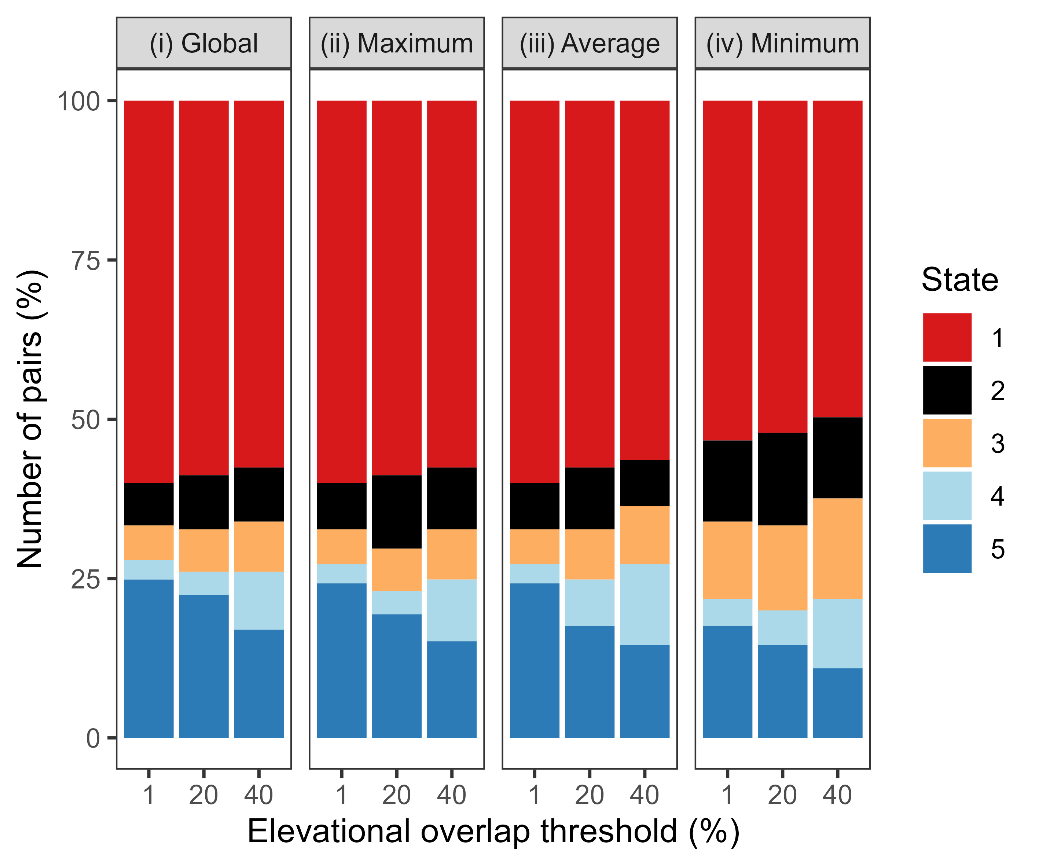

**Figure S6** Distribution of sister pairs among states for different elevational overlap criteria. Colours corresponds to the five states in Figure 1, and the proportion of sister pairs assigned to this state is indicated by bar height. The four columns represent the four protocols used to assign sister pairs to the states, and the x-axis shows the threshold of overlap between elevational ranges of sister species below which sister species are considered elevationally non-overlapping. For further description of protocols and impact of state assignment see Figure S3-S5. The results in the main text are for the ‘average overlap’ protocol using the 20% elevational overlap threshold.

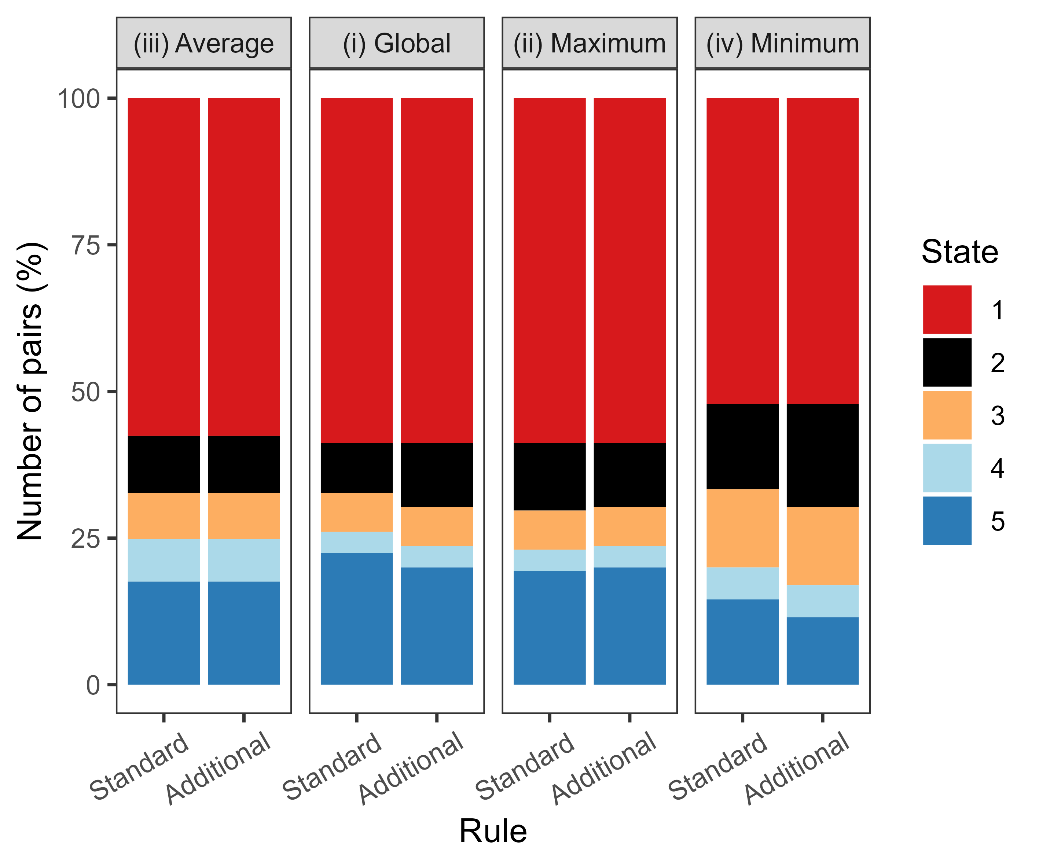

**Figure S7** Assignment of sister pairs to states according to different overlap protocols. Colours corresponds to the five states in Figure 1, and the proportion of sister pairs assigned to this state is indicated by bar height. The four columns represent the four protocols used to assign sister pairs to the states, and the x-axis shows the two scenarios under which sister pairs are assigned to state 2: ‘standard’ referring to assignment based on <20% overlap in elevation in sympatry and ≥20% in allopatry, and ‘additional’ under which a 20% decrease in elevational overlap from allopatry to sympatry is used. All other states are assigned based on a 20% elevational overlap criterion. For further description on protocols see Figure S3. The results in the main text are for the ‘average overlap’ protocol using the 20% elevational overlap threshold and ‘standard’ rule.

*Accuracy and reliability*

In order to assess the accuracy and precision of the model in recovering the transition rates we assessed five separate scenarios (S1-S5). Scenarios S1 to S3 refer to respectively cases where >80% of the sister pairs that leave state 1 (i.e. allopatry with overlapping elevation ranges) go through either elevational (S1) sorting (1->3->5), (S2) displacement (1->2->5 or 1->2->4->5) or (S3) conservatism (1->5). These scenarios are arbitrary but were chosen to be both realistic (in terms of the order-of-magnitude variation in rates observed) while allowing us to test for any bias to over or underestimate the prevalence of a particular community assembly pathway when it dominates the empirical data. For S4, we model the opposite case, where transition rates between all states are equal (0.05) to investigate if there is any bias in overestimating any particular rate. We chose a transition rate of 0.05 because it is comparable to the initial rates (i.e. r12, r13, r15) estimated for the empirical data. Using a higher value (e.g. 0.25), would be expected to lead to more accurate and precise estimates for the latter rates (e.g. r24 and r25) because of the larger number of sister pairs in these states, but would not be reflective of the initial rates that were estimated. Finally, for S5 we test how well we can recover the true rates from a scenario where we simulate under the empirical model estimates. While this is not an exhaustive search of model performance and additional parameter combinations and values could be explored, the scenarios were chosen to bracket a range of limiting but realistic scenarios, assuming either equivalence among or dominance of a particular community assembly pathway.

**Table S1** Estimates of simulated transition rates derived from the model-averaged approach. For each scenario 100 datasets were simulated forwards in time under the true rates using the observed sister pair ages. Species pairs could transition between states until the simulation reached the pair’s age. Subsequently, Markov models were fitted to the simulated data to estimate transition rates (r12, r13, r15, r24, r25, r34, r45,). The first three scenarios (S1, S2, & S3) represent scenarios where >80% of pairs go through elevational sorting, displacement and conservatism respectively. Under scenario S4 rates are equal among transitions and in S5 transition rates used in the simulation are the rates estimated for the empirical data. For each dataset model-averaged parameter values were obtained by AIC weighted-averaging of the estimated parameters for all models that were within 2 AIC of the best model. Median and the 95%CI (within parentheses), were constructed per scenario from the model-averaged estimates across the estimated parameters for the 100 datasets. Rate coverage was calculated by counting the *n* cases out of 100 in which the true rates were within AIC-weighted 95%CIs. The CIs were constructed for every dataset based on the set of models that were within 2 AIC of the best model.

|  | **S1 - Elevational sorting** | | |  | **S2 - Elevational displacement** | | |
| --- | --- | --- | --- | --- | --- | --- | --- |
|  | **True rate** | **Estimated** | **Coverage** |  | **True rate** | **Estimated** | **Coverage** |
| **r12** | 0.01 | 0.01 (0-0.03) | 98 |  | 0.1 | 0.09 (0.07-0.14) | 93 |
| **r13** | 0.1 | 0.09 (0.06-0.13) | 93 |  | 0.01 | 0.01 (0-0.02) | 92 |
| **r15** | 0.01 | 0.01 (0-0.02) | 85 |  | 0.01 | 0.01 (0-0.03) | 92 |
| **r24** | 0.01 | 0.07 (0.02-0.58) | 0 |  | 0.1 | 0.09 (0.02-0.23) | 86 |
| **r25** | 0.01 | 0.05 (0.02-0.15) | 0 |  | 0.1 | 0.07 (0.01-0.13) | 73 |
| **r34** | 0.1 | 0.09 (0.02-0.18) | 84 |  | 0.01 | 0.07 (0.02-0.49) | 1 |
| **r45** | 0.1 | 0.06 (0.02-0.33) | 54 |  | 0.01 | 0.06 (0.02-0.4) | 2 |
|  | **S3 - Elevational conservatism** | | |  | **S4 – Equal rates** | | |
|  | **True rate** | **Estimated** | **Coverage** |  | **True rate** | **Estimated** | **Coverage** |
| **r12** | 0.01 | 0.01 (0.01-0.06) | 67 |  | 0.05 | 0.05 (0.03-0.08) | 98 |
| **r13** | 0.01 | 0.01 (0-0.03) | 88 |  | 0.05 | 0.05 (0.03-0.07) | 97 |
| **r15** | 0.1 | 0.09 (0.05-0.13) | 95 |  | 0.05 | 0.05 (0.02-0.07) | 97 |
| **r24** | 0.01 | 0.07 (0.03-2.09) | 2 |  | 0.05 | 0.05 (0.03-0.17) | 89 |
| **r25** | 0.01 | 0.06 (0.03-1.87) | 1 |  | 0.05 | 0.05 (0.03-0.29) | 84 |
| **r34** | 0.01 | 0.08 (0.03-1.27) | 7 |  | 0.05 | 0.05 (0.03-0.22) | 92 |
| **r45** | 0.01 | 0.06 (0.03-4.2) | 0 |  | 0.05 | 0.06 (0.04-0.86) | 57 |
|  | **S5 – Estimated rates** | | |  |  |  |  |
|  | **True rate** | **Estimated** | **Coverage** |  |  |  |  |
| **r12** | 0.08 | 0.08 (0.06-0.14) | 99 |  |  |  |  |
| **r13** | 0.06 | 0.07 (0.03-0.1) | 97 |  |  |  |  |
| **r15** | 0.08 | 0.07 (0.04-0.1) | 92 |  |  |  |  |
| **r24** | 0.17 | 0.12 (0.06-0.36) | 78 |  |  |  |  |
| **r25** | 0.06 | 0.09 (0.05-0.34) | 71 |  |  |  |  |
| **r34** | 0.18 | 0.19 (0.06-0.59) | 83 |  |  |  |  |
| **r45** | 0.08 | 0.09 (0.05-0.64) | 70 |  |  |  |  |

**Table S2** Estimates of simulated transition rates derived from the ‘best’ model approach. For each of the 100 datasets per scenario this table represents the parameters estimated by the ‘best’ model. Median and the 95%CI, between parentheses, were calculated per scenario from the ‘best’ model estimates across these datasets. Additionally, rate coverage is shown referring to the *n* cases out of 100 in which the confidence intervals of the estimates for the individual ‘best’ model capture the true rates. For details see Table S1.

|  | **S1 - Elevational sorting** | | |  | **S2 - Elevational displacement** | | |
| --- | --- | --- | --- | --- | --- | --- | --- |
|  | **True rate** | **Estimated** | **Coverage** |  | **True rate** | **Estimated** | **Coverage** |
| **r12** | 0.01 | 0.01 (0-0.04) | 78 |  | 0.1 | 0.1 (0.06-0.14) | 89 |
| **r13** | 0.1 | 0.09 (0.06-0.12) | 94 |  | 0.01 | 0.01 (0-0.02) | 88 |
| **r15** | 0.01 | 0.01 (0-0.02) | 91 |  | 0.01 | 0.01 (0-0.03) | 81 |
| **r24** | 0.01 | 0.04 (0-0.94) | 47 |  | 0.1 | 0.09 (0.01-0.23) | 69 |
| **r25** | 0.01 | 0.02 (0-0.35) | 51 |  | 0.1 | 0.09 (0-0.15) | 55 |
| **r34** | 0.1 | 0.09 (0-0.21) | 77 |  | 0.01 | 0.07 (0-0.87) | 42 |
| **r45** | 0.1 | 0.06 (0-0.54) | 45 |  | 0.01 | 0.02 (0-1.12) | 50 |
|  | **S3 - Elevational conservatism** | | |  | **S4 – Equal rates** | | |
|  | **True rate** | **Estimated** | **Coverage** |  | **True rate** | **Estimated** | **Coverage** |
| **r12** | 0.01 | 0.01 (0-0.08) | 79 |  | 0.05 | 0.05 (0.04-0.08) | 88 |
| **r13** | 0.01 | 0.01 (0-0.04) | 87 |  | 0.05 | 0.05 (0.03-0.07) | 89 |
| **r15** | 0.1 | 0.09 (0.01-0.13) | 85 |  | 0.05 | 0.05 (0.02-0.07) | 93 |
| **r24** | 0.01 | 0.08 (0.01-0.68) | 25 |  | 0.05 | 0.05 (0.02-0.19) | 83 |
| **r25** | 0.01 | 0.02 (0.01-3.34) | 51 |  | 0.05 | 0.05 (0.02-0.41) | 83 |
| **r34** | 0.01 | 0.09 (0.01-1.01) | 28 |  | 0.05 | 0.05 (0.02-0.41) | 83 |
| **r45** | 0.01 | 0.01 (0.01-4.54) | 60 |  | 0.05 | 0.05 (0.02-1.24) | 79 |
|  | **S5 – Estimated rates** | | |  |  |  |  |
|  | **True rate** | **Estimated** | **Coverage** |  |  |  |  |
| **r12** | 0.08 | 0.08 (0.05-0.16) | 74 |  |  |  |  |
| **r13** | 0.06 | 0.07 (0.03-0.11) | 74 |  |  |  |  |
| **r15** | 0.08 | 0.07 (0.02-0.11) | 81 |  |  |  |  |
| **r24** | 0.17 | 0.11 (0.04-0.42) | 36 |  |  |  |  |
| **r25** | 0.06 | 0.08 (0.01-0.41) | 46 |  |  |  |  |
| **r34** | 0.18 | 0.2 (0.02-0.6) | 50 |  |  |  |  |
| **r45** | 0.08 | 0.08 (0.01-0.6) | 54 |  |  |  |  |

*How often do we correctly infer the prevalence of community assembly pathways?*

To estimate the frequency at which we correctly infer the prevalence of each community assembly pathway (i.e. ES, ED, or EC) we again follow a multi-step approach. First, with the rates recovered for the empirical data (e.g. Table S1 scenario 5), we simulate 100 primary datasets for which we know the states that each sister pair will have passed through. Second, for each dataset we perform the same modelling procedures as for the empirical data to estimate the transition rates – note step 1 and 2 are identical to recovering the rate estimates for Table S1. Third, using the rates estimated per dataset, we simulate 100 secondary datasets per primary dataset, totalling 10000 datasets. During the simulation of these secondary datasets we again take note of the states through which each sister pair passes. Fourth, from the prevalence of each pathway in the secondary datasets we construct 95% confidence intervals for the pathway of interest (Figure 2b, c, d). Finally, we evaluated if the prevalence of the pathways in the primary dataset falls within the 95%CI of the respective secondary datasets. We term the number of times that the 95%CI capture the prevalence of the primary dataset ‘pathway coverage’.

**Table S3** recovery of pathways taken according to the best models for each primary dataset. The procedure in which results are obtained are described above. Figure refers to the pathways in Figure 2: (b) sister pairs that have left state 1 and transitioned to state 3, 2, or 5 representing elevational sorting (ES), displacement (ED), or conservatism (EC), (c) sister pairs that occupy non-overlapping elevational ranges in sympatry that have arisen through the ES or ED pathway, (d) sister pairs that are sympatric with overlapping elevational ranges that have arisen through ES, ED, or EC. Pathway coverage refers to the % of times the true prevalence of each pathway as simulated in the primary dataset falls within the 95%CI’s of the secondary dataset. For primary prevalences that fall outside of the 95%CI, the distance to the mean prevalence under the secondary simulations is expressed in standard deviations.

| **Figure** | **Process** | **Pathway coverage** | **Under- estimated** | **Over- estimated** | **Dist. to mean** | **SDs to mean** |
| --- | --- | --- | --- | --- | --- | --- |
| 2b | ES | 85 | 3 | 12 | 0.15 | 2.74 |
| 2b | ED | 86 | 2 | 12 | 0.23 | 4.26 |
| 2b | EC | 80 | 20 | 0 | 0.2 | 27.65 |
| 2c | ES | 80 | 9 | 11 | 0.21 | 3.72 |
| 2c | ED | 80 | 11 | 9 | 0.21 | 3.72 |
| 2d | ES | 84 | 5 | 11 | 0.21 | 10.83 |
| 2d | ED | 68 | 6 | 26 | 0.33 | 22.24 |
| 2d | EC | 58 | 36 | 6 | 0.33 | 17.49 |

**Table S4** recovery of pathways taken according to the model-averaged parameters for each primary dataset. See Table S3 for details.

| **Figure** | **Process** | **Pathway coverage** | **Under- estimated** | **Over- estimated** | **Dist. to mean** | **SDs to mean** |
| --- | --- | --- | --- | --- | --- | --- |
| 2b | ES | 94 | 1 | 5 | 0.13 | 2.28 |
| 2b | ED | 87 | 2 | 11 | 0.18 | 3.27 |
| 2b | EC | 83 | 17 | 0 | 0.15 | 3.12 |
| 2c | ES | 88 | 6 | 6 | 0.21 | 3.28 |
| 2c | ED | 88 | 6 | 6 | 0.21 | 3.28 |
| 2d | ES | 92 | 3 | 5 | 0.19 | 3.45 |
| 2d | ED | 72 | 3 | 25 | 0.26 | 4.46 |
| 2d | EC | 70 | 29 | 1 | 0.29 | 3.39 |

*Initial state analysis*

In the main text we made the assumption that at the time of divergence, sister species will occur in allopatry and have overlapping elevational ranges (state 1). That almost all (>95% of) speciation events in birds involve a stage of geographic isolation is well established (Phillimore *et al.*, 2008; Price, 2008). However, it is possible that newly formed allopatric species may also arise with differentiated elevational ranges (i.e. state 3) if for example the ancestral species exhibits regional variation in its elevation range. To explore this possibility and how this may influence our conclusions we conducted a further analysis in which we included an additional parameter in our model, *γ*, which is constrained to occur from 0 to 1 and describes the probability that the initial state is state 3. Thus, 1- *γ* is the probability that the initial state is state 1.

We used maximum likelihood (ML) to simultaneously estimate the transition rates and *γ.* We also estimated a likelihood profile by incrementally varying *γ* in steps of 0.01 from 0 to 0.3 in order to understand the shape of the likelihood surface and identify values of *γ* within the 95%CI of the ML estimate (i.e. within 1.92 log-likelihood units). Preliminary analyses of larger step sizes showed that the likelihood profile did not have multiple peaks, supporting our choice of step sizes. We applied both of these approaches to our original transition matrix which includes the following transitions (r12, r13, r15, r24, r25, r34, r45). We estimate *γ* = 9.80e-15 (95%CI: 0-0.08), indicating that few if any sister pairs are estimated to arise in state 3 (Figure S8a). This model assumes that sister pairs arising in state 1 can transition to state 3 (in which sister pairs can also arise with probability *γ*). However, it does not consider the possibility that sister pairs arising in state 3 can transition to state 1 (i.e. they converge in their elevation ranges in allopatry). We therefore repeated our analysis using a final scenario where we instead allow transitions from state 3 to state 1 but not the reverse (i.e. r12, r15, r24, r25, r31, r34, r45). Note we do not allow two-way transitions between state 1 and 3 as rates would be unidentifiable. In this model, the proportion of sister pairs estimated to arise in state 3 is not zero (*γ* = 0.079 (95%CI 0.05-0.13) but is very low compared to the proportion that arise in state 1 (Figure S8b). We note that the ML estimate of the latter scenario is >1.92 log-likelihood units below the 95%CI of the former, indicating that the model with transitions from state 3 to 1 is a worse fit.

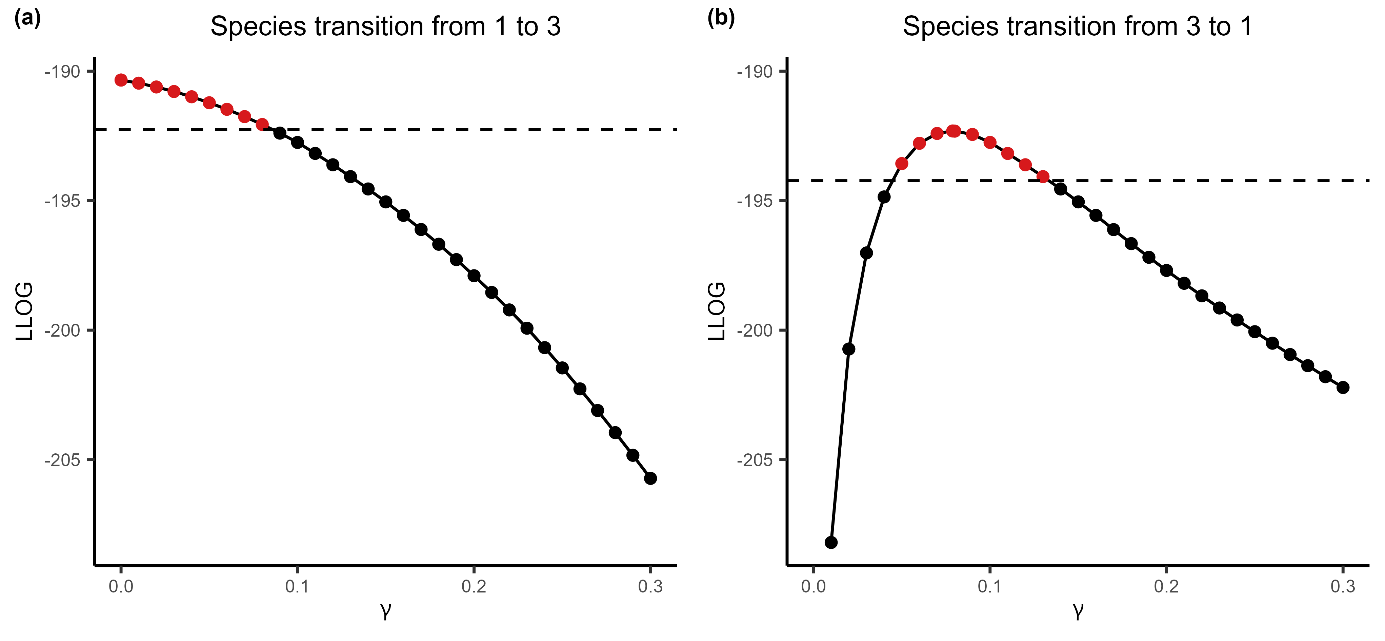

**Figure S8** Likelihood of elevational differentiation upon speciation (*γ*) i.e. that state 3 is the initial state. (a) shows the likelihood profile for models where sister species can transition from allopatry without elevational differentiation to allopatry with elevational differentiation (state 1 → 3), and (b) shows the profile for models with the opposite transition (state 3 → 1) (Figure 1). Values of *γ* above the dashed line (95%CI) are highlighted in red and considered not significantly different from the maximum likelihood estimate.

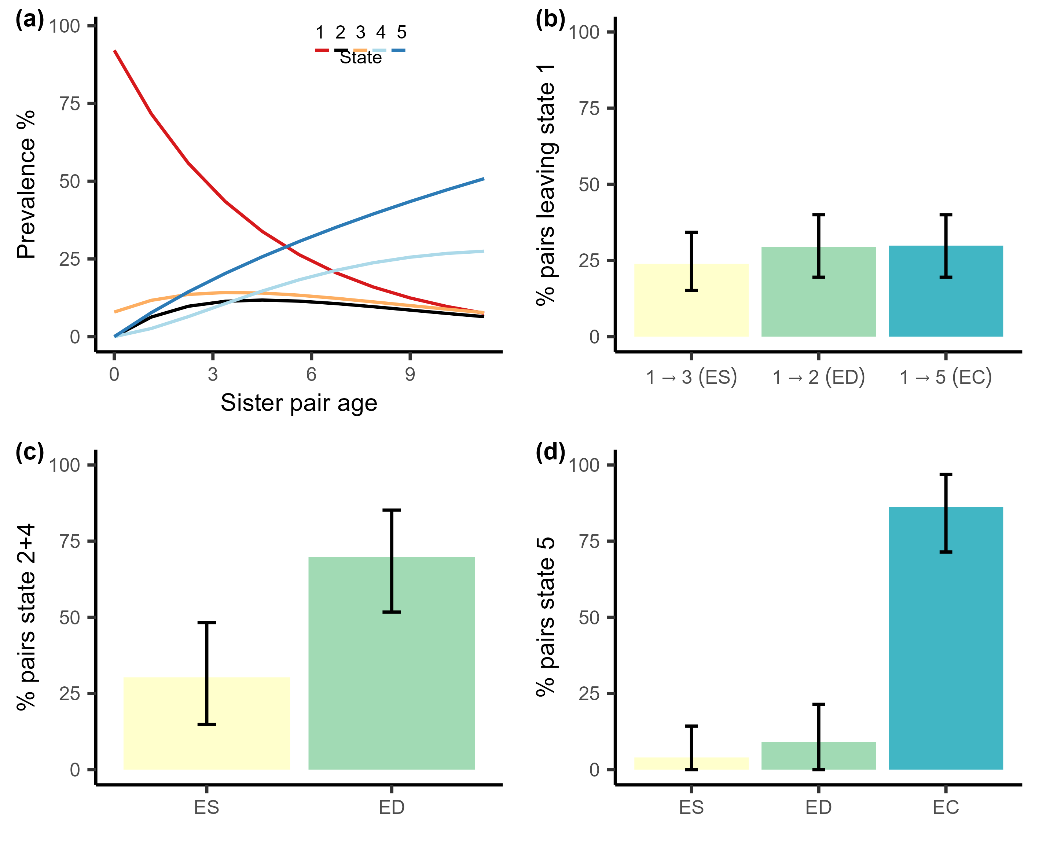

**Figure S9** The inferred community assembly pathways of sister species when species can arise in allopatry with non-overlapping elevations (i.e. state 3). a) estimated prevalence of the five states with time since divergence, as represented in Figure 1. The lines show the average prevalence across 1000 simulations. b) Percentage of sister pairs that have left state 1 and have transitioned to state 3, 2, or 5 representing the elevational sorting (ES), displacement (ED), or conservatism (EC) pathways respectively. c) the percentage of pairs that currently occupy non-overlapping elevational ranges in sympatry (i.e. state 2 or 4) that have arisen through the ES or ED pathway. d) The percentage of pairs that are sympatric with overlapping elevational distributions (i.e. state 5) that have arisen through the ES, ED, or EC pathway. Results are obtained from posterior-predictive simulations of the model-averaged parameter estimates for an elevational differentiation threshold of 20% according to the ‘average’ elevation range overlap protocol, and 8% chance of arising in state 3.

*Calculating the lag time to sympatry*

The rate of range expansions leading to sympatry may be accelerated by the capacity for species to occur on the same slope without having to first diverge (and then expand) in their elevation range i.e. the elevation conservatism (EC) pathway. To determine the extent to which EC shortens the lag time to sympatry, we compared the mean waiting time to sympatry inferred by our models, either (1) assuming this pathway is present or (2) this pathway is absent. To do this we simulated the dynamics of *n* sister species pairs (*n* = 100k) forward in time using a Gillespie algorithm. Pairs transitioned between states according to the transition rates recovered under the model-averaged rates for the ‘average overlap’ protocol using a 20% elevational overlap threshold. Simulations were run until all sister species had reached the final state 5 – sympatry with overlapping elevational ranges. For each sister pair we tracked the first point in time at which sympatry is attained, i.e. state 2 for ED, state 4 for ES, and state 5 for EC (Figure 1). We calculated the expected lag time to sympatry by averaging lag times across sister pairs. We then repeated this procedure but setting the transition rate from state 1 to 5 to zero, and thus preventing species from using the EC pathway. The comparison of lag times under these two models thus indicates the extent to which the ability of species to attain sympatry without diverging in their elevation ranges (i.e. EC) accelerates the attainment of sympatry following speciation.
